# SERM: a self-consistent deep learning solution for rapid and accurate gene expression recovery

**DOI:** 10.1101/2022.01.18.476789

**Authors:** Md Tauhidul Islam, Jen-Yeu Wang, Hongyi Ren, Xiaomeng Li, Masoud Badiei Khuzani, Shengtian Sang, Lequan Yu, Liyue Shen, Wei Zhao, Lei Xing

## Abstract

Single cell RNA sequencing (scRNA-seq) is a promising technique to determine the states of individual cells and classify novel cell subtypes. Computationally, the processing of scRNA-seq data presents a daunting challenge because of the noisy nature and humongous size and dimensionality of the data. Compromised solution by omitting the genes with low expression is commonly taken in current scRNA-seq analysis, which leads to inaccurate gene counts. In this paper, we introduce a broadly applicable data-driven gene expression recovery framework, referred to as the self-consistent expression recovery machine (SERM), to impute the missing gene expression. Using deep learning, SERM first learns from a subset of the noisy gene expression data to estimate the underlying data distribution. SERM then recovers the overall gene expression data by imposing a self-consistency on the gene expression matrix, thus ensuring that the expression levels are similarly distributed in different parts of the matrix. We show that SERM significantly improves the accuracy of gene imputation with at least 100-fold increase in computational efficiency in comparison to the state-of-the-art techniques. Thus SERM promises to provide an urgently needed computational solution for rapid and accurate recovery of big genomic expression data. SERM is available as a web-based computational tool (https://www.analyxus.com/compute/serm) and its source codes can be found in https://github.com/xinglab-ai/self-consistent-expression-recovery-machine.

## Introduction

Single-cell RNA sequencing (scRNA-seq) has emerged as an effective tool for variety of cellular analysis tasks such as quantifying the state of individual cells^1,2^, classifying novel cell subtypes^3,4^, assessing progressive gene expression (cell trajectory analysis)^5–7^, performing spatial mapping^8,9^ and identifying differentially expressed genes^10,11^. Despite its prevalence in computational biology, due to noise and low transcript capture efficiency, the resulting gene expression matrix from scRNA-seq is typically sparse, which often results in a loss of important biological information^12–14^. In the past decade, intense research has been devoted to address the computational challenges in recovering the omitted gene expressions. The developed techniques can be broadly divided into three categories^15^. The first category models the sparsity of expression data using probabilistic models. SAVER^12^, SAVER-X^16^, bayNorm^17^, scImpute^13^, scRecover^18^ and VIPER^19^ are the prominent methods of this group. The second group are the imputation methods that utilize averaging/smoothing to recover the expression values. DrImpute^20^, MAGIC^14^ and k-NN smoothing^21^ are the most popular techniques of this category. The last group of techniques are based on reconstruction of data either using deep learning (AutoImpute^22^, DCA^23^, DeepImpute^24^, SAUCIE^25^, scScope^26^, scVI^27^) or low-rank matrix assumption (mcImpute^28^, PBLR^29^). Overall, the model-based techniques assume a specific model for the expression data, which may limit their application in some practical cases. As an example, SAVER extracts information from correlated genes and employs the penalized regression models to impute the values. It assumes that the gene expression count of a cell follows a gamma-Poisson distribution, which may not be appropriate in many cells, including those with low gene expression values^12^. The smoothing-based methods extract information from similar cells to impute the expression data. However, the averaging effect in these methods may potentially eliminate variability in gene expressions across cells^14^. The deep-learning based methods learn from the data to perform imputation and do not impose any assumption. However, as the deep learning techniques are purely data-driven without incorporation of any physical model, the data imputation becomes complex black-box operations and thus difficult to control. As a result, the obtained results may be distorted and lack trustworthiness. Low-rank matrix based techniques such as mcImpute use a matrix completion approach to impute the data with considerations of both gene-gene and cell-cell relationships. However in mcImpute, all zero expression values are treated as drop out events, which may cause spurious results in many practical scenarios. From the perspective of computational efficiency, the analytical techniques (model-based, smoothing-based and low rank matrix-based methods) are in general not scalable to large datasets since the entire dataset must be processed as a whole.

Here we propose the self-consistent expression recovery machine (SERM) that employs a data-driven strategy to recover the gene expression data. Computationally, the SERM first learns the underlying data distribution using deep learning and then utilizes self-consistency of the data to recover the actual expression data. Specifically, we leverage a deep learning optimization to extract the latent representation of the data and reconstruct the denoised expression values (Fig. 1-step 1). The SERM then uses a curve fitting technique to learn a probability distribution from a parametric family to represent the reconstructed expression values from the optimization (Fig. 1-steps 2, 3). Next, a rectangular region of interest (ROI) is selected within the expression matrix (Fig. 1-step 4) and imputation is performed based on histogram equalization using the learned probability distribution (Fig. 1-step 5). The imputation calculation proceeds iteratively, with the ROI shifting along the x- and y-axes of the gene expression matrix for each new iteration. This iterative process can be regarded as a ‘sliding window’ approach (Fig. 1-step 6). In the last step, all the ROIs are interpolated using bilinear resampling to achieve the global consistency of the expression values (Fig. 1-step 7, supplementary section 1). Because of this divide-and-conquer strategy, SERM is scalable to large datasets and can impute the datasets much faster than other analytical methods as well as most of the deep-learning based methods.

**Figure 1.**
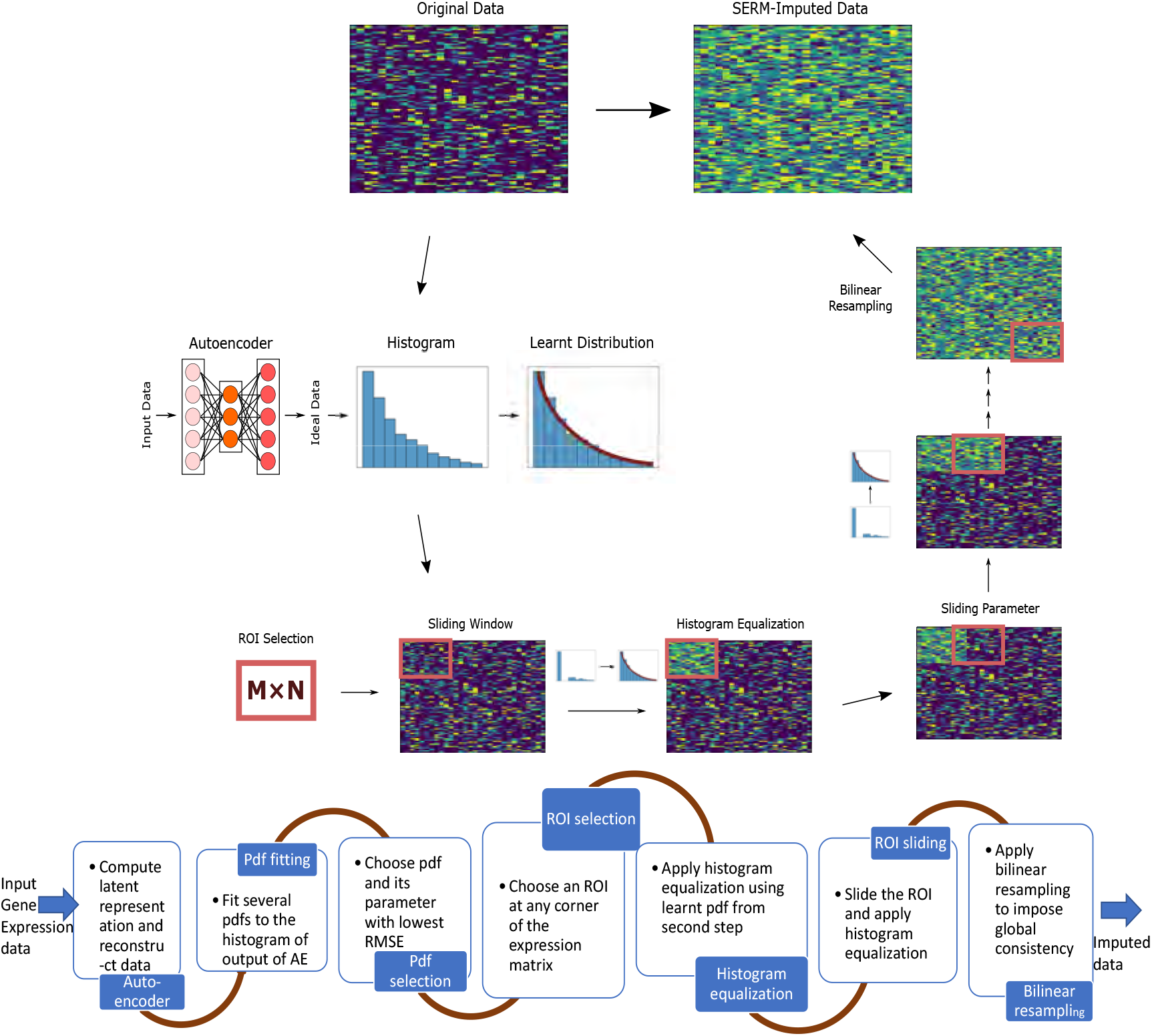
Workflow of SERM. In SERM, at first a subset of the noisy expression matrix is given input to an encoder-decoder network. The distribution function that best describes the reconstructed data by the decoder is learned using fitting of different pdfs. In the next step of SERM, an ROI is selected and histogram equalization is performed on that ROI using the learned pdf in the previous step. The ROI is then slided along x and y direction throughout the whole expression matrix and histogram equalization is performed on each ROI. All the regions are then interpolated using bilinear resampling in the final step to impose the global consistency.

We showcase the efficacy and accuracy of SERM on various benchmark datasets and demonstrate its superior performance over the existing baseline methods. Compared to alternative imputation techniques, SERM consistently achieves significantly better imputation accuracy, and speed. The technique promises to substantially improve the way that ScRNAseq is processed and its applications in biomedical research.

## Results

In this section, we first present the synthetic and benchmark gene sequence datasets to demonstrate SERM’s efficacy in visualization and clustering. We then examine SERM’s performance by analyzing cell trajectory datasets. Throughout this study, we describe how SERM can be used to achieve higher classification accuracy and better visualization. We also demonstrate how SERM can be used to impute very big genomic data, namely the human cell landscape and the mouse cell atlas, to discover novel biological information.

In our following analyses, for cluster and visualization datasets, we judge the performance of six imputation methods (MAGIC, mcImpute, AutoImpute, DeepImpute, SAUCIE and SERM) in terms of 1) visualization quality, 2) correlation of the imputed data with original data, 3) clustering accuracy, 4) normalized mutual information (NMI) and 4) three cluster quality indices (adjsted Rand (AR), Rand and Hubert). For trajectory analysis datasets, the performance of different methods is judged based on 1) trajectory quality, 2) correlation of the imputed data with original data and 3) preservation of time point to time point distance.

### SERM provides better data recovery for superior visualization and clustering

For our first example, we use the Splatter simulator to generate simulated data with 20 classes^30^. Each class has 1,000 cells and 5,000 genes. Classes 1-20 have the same rate parameter of 0.9 and shape parameters of 0.10, 0.11, 0.15, 0.16, 0.20, 0.21, 0.25, 0.26, 0.30, 0.31, 0.35, 0.36, 0.40, 0.41, 0.50, 0.52, 0.70, 0.71, 0.80 and 0.805, respectively. Dropout shape is set to −1, dropout midpoint is set to zero and dropout type is set to experiment in the Splatter simulation. Other simulation parameters are set at default numbers.

The first two principal components and the global histogram of the dataset are shown in Fig. 2 (a1) and (b1). Due to a high dropout rate, we observe that the principal components are not able to differentiate among the data classes. The histogram has a very large number of data points in the zero bin, whereas there are very small numbers of points in the other non-zero bins. The principal components from MAGIC- and mcImpute-imputed data are shown in (c1) and (a2). We observe degradation of the cluster quality in the principal components. The histograms are shown in (d1) and (b2), where it is seen that MAGIC changes the data mean along with the distribution, resulting in data distortion. SERM, on the other hand, distributes the zero values to non-zero values to preserve the learned exponential distribution of the data values, which results in a superior data representation by principal components (c2) and a superior histogram (d2). The distributions of ideal data values are shown in Fig. S1, which resemble the distribution from SERM-imputed data better than that from other methods. The Pearson coefficient between the dropout-free data with unimputed data and imputed data from different methods (MAGIC, mcImpute, AutoImpute, DeepImpute, SAUCIE and SERM) are shown in Fig. 2 (a3)-(d3) and Fig. S2. The unimputed, MAGIC-imputed and SAUCIE-imputed data show poor Pearson coefficient. McImpute and DeepImpute performs comparatively better than other methods in this case. The SERM-imputed data results in the highest Pearson coefficient, which proves that it has the best match with the dropout-free data.

**Figure 2.**
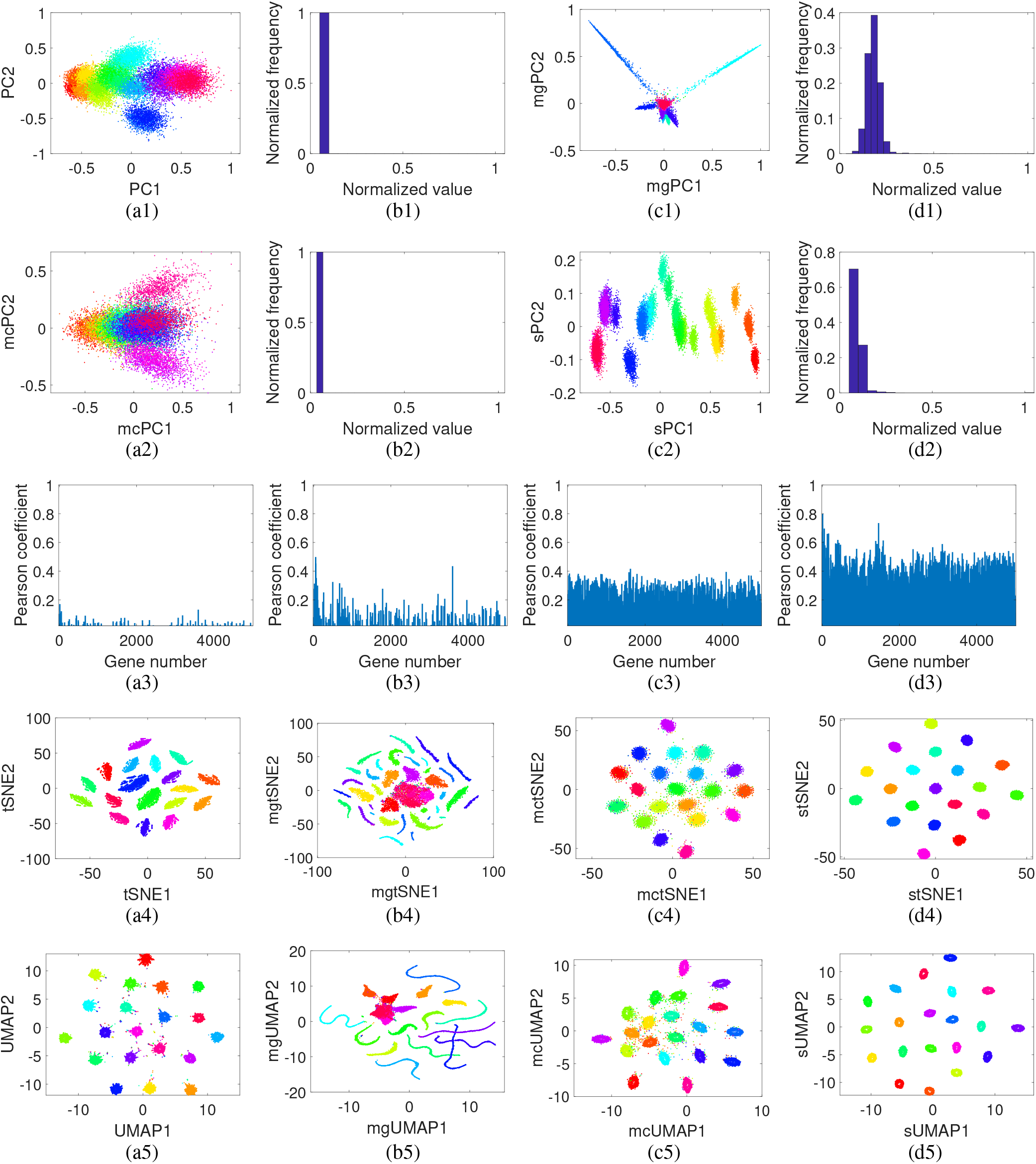
(a1)-(d5) Genomic data with large number (20) of classes simulated with Splatter simulator. Classes 1-20 have the same rate parameter of 0.9 and shape parameters of 0.10, 0.11, 0.15, 0.16, 0.20, 0.21, 0.25, 0.26, 0.30, 0.31, 0.35, 0.36, 0.40, 0.41, 0.50, 0.52, 0.70, 0.71, 0.80 and 0.805, respectively. Dropout shape is set to 1, dropout midpoint is set to zero and dropout type is set to experiment in Splatter simulation. Other simulation parameters are set at default numbers. The first two principal components of the original simulated data are shown in (a1) and its histogram in (b1). The first two principal components of MAGIC, mcImpute and SERM imputed data are shown in (c1), (a2) and (c2) and their histograms in (d1), (b2) and (d2), respectively. The learned distribution by SERM in this case is exponential with parameter value of 20. The principal components of SERM imputed data clearly show the data classes, whereas that from other methods fail to do so. Correlation coefficient between the gene expressions of the unimputed (a3)/imputed data and dropout-free data for MAGIC (b3), mcImpute (c3) and SERM (d3). Visualization of unimputed and imputed data by t-SNE and UMAP. t-SNE results from original data and imputed data from MAGIC, mcImpute and SERM are shown in (a4)-(d4) and UMAP results from them are shown in (a5)-(d5), respectively. t-SNE and UMAP results from SERM imputed data are much better in separating the classes, whereas MAGIC degrades the data as a result of imputation.

The superior visualization and classification from SERM-imputed data become more obvious when we see the t-SNE and UMAP results from data imputed by different methods, as shown in Fig. 2 (a4)-(d5) and Fig. S2. As a result of the dropout effect, the t-SNE and UMAP visualizations of the original data show data classes with very small distances and many randomly clustered data points (a4, a5). McImpute partially succeeds in imputing the data, which is reflected in the visualizations (c4, c5). MAGIC, AutoImpute and SAUCIE, on the other hand, distorts the data visualization (Fig. 2 (b4, b5) and Fig. S2 (a1-a3, c1-c3). DeepImpute performs better than all other techniques except SERM. SERM successfully imputes the data and preserves the quality of visualization, which can be seen in Fig. 2 (d4, d5).

In the next study, we use benchmark data from four different experiments (cellular taxonomy of the bone marrow stroma in Homeostasis and leukemia^31^, classification of cells from mammalian brains^32^, data from mouse intestinal epithelium^33^ and comparison of engineered 3D neural tissues^34^) available from the the single cell database from the Broad Institute. Descriptions regarding how the data was acquired in these experiments can be found in the methods section.

In this analysis, a percentage of the original data is filled with zeros and different methods are then used to recover the original data. We show t-SNE maps of all four datasets (with 20% of the data filled with zeros) imputed by different techniques in Fig. 3 and Fig. S3. The t-SNE maps of the original data are shown Fig. 3 (a1, a2, a3, a4). The data classes are better separated in SERM-imputed data as shown in Fig. 3 (d1-d4) similar to the original data. MAGIC and SAUCIE distort the data visualizations in most cases (b1, b2, b3). McImpute and DeepImpute fail to provide imputation as good as SERM in all cases. AutoImpute performs comparatively in terms of visualization quality to SERM and better than other methods.

**Figure 3.**
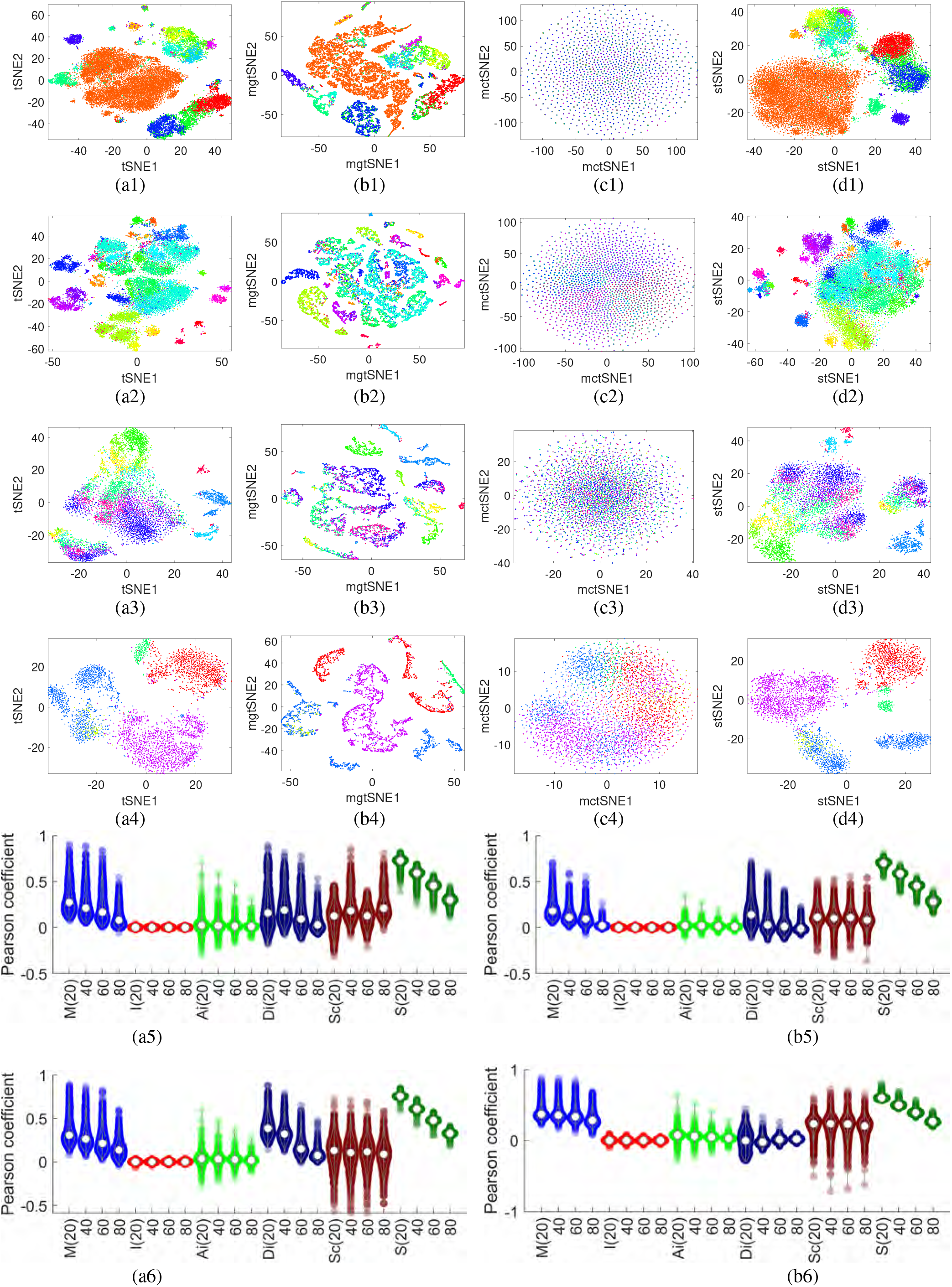
(preceding page). t-SNE results from original data (a), imputed data from MAGIC (b), mcImpute (c) and SERM (d) for (1) cellular taxonomy, (2) mammalian brain, (3) mouse intestinal epithelium and (4) 3D neural tissue data. All the classes are better visualized in results from SERM imputation. MAGIC and mcImpute distort the data in many cases (b1, b2, b3, b4, c4), whereas SERM keeps the consistency of the data intact in every cases. Legends are added in the supplementary section 2. Pearson coefficient between the imputed data and ground truth data for all the methods are shown for cellular taxonomy (a5), mammalian brain (b5), mouse intestinal epithelium (a6) and 3D-effect data (b6), respectively. In (a5-b6), ‘M’ denotes MAGIC, ‘I’ denotes mcImpute, ‘AI’ denotes AutoImpute, ‘DI’ denotes DeepImpute, ‘Sc’ denotes SAUCIE and ‘S’ denotes SERM methods, respectively. The percent of randomly-zeroed expression values (20, 40, 60 and 80) in the ground truth data to create the noisy data are shown in braces. The spread of the violin denotes the variance of the coefficient values across different cells.

The quantitative results for the imputed data from different methods are shown in Fig. 3 (a5-b6) and Fig. S8 (a-d). When 20% of the data are filled with zeros, it is seen that SERM provides a Pearson coefficient of more than 0.7 in most cases, whereas all other methods provide less than 0.4. DeepImpute and MAGIC perform comparatively better than other techniques. Even when 80% of the data are filled with zeros, SERM recovers the original expression data with 0.2-0.4 Pearson coefficient, whereas the other methods show very poor coefficient (≈ 0). Moreover, the small variances of Pearson coefficient found in case of SERM-imputed data proves the reliability and consistency of its high accuracy in recovering the true expression values. Similar to Pearson coefficient, SERM excels in all other quantitative indices (AR, Rand, accuracy, NMI and Hubert) as shown in Fig. S8 (a-d).

### SERM offers distortionless cell trajectory analysis

The aim of the following experiments is to demonstrate distortionless and accurate cell trajectory analysis enabled by SERM. Trajectory mapping entails preprocessing the raw data; failing to do so yields distorted results. As an example, recently developed dimensionality reduction method PHATE^35^ has been shown to accurately infer cell trajectory maps. However, performance of this method (and all other trajectory inference methods) can be degraded when biological information is obscured by technical noise such as high dropout. SERM reduces the effect of dropout and thus can assist PHATE (and all other trajectory inference methods) in accurately analyzing cell trajectory.

The first dataset we consider is acquired by profiling 38,731 cells from 694 embryos across 12 closely separated stages of early zebrafish development^36^ using a massively parallel scRNA-seq technology named Drop-seq^2^. Data were acquired from the high blastula stage (3.3 hours postfertilization (hpf), moments after transcription starts from the zygotic genome) to the six-somite stage (12 hours after postfertilization, just after gastrulation). Most cells are pluripotent at the high blastula stage, whereas many cells have differentiated into specific cell types at the six-somite stage. Similar to the cluster data analysis, a percentage of the original data is filled with zeros and different methods are then used to recover the original data.

PHATE visualizations of original data and data imputed by all six methods are shown in Fig. 4 (a1-d1) and Fig. S9. It is observed from the figure that only AutoImpute and SERM-imputed data produce results with less distortion. Results from imputed data by other methods are highly distorted, and it is difficult to find any trajectory. DeepImpute performs comparatively better than SAUCIE and MAGIC in this case.

**Figure 4.**
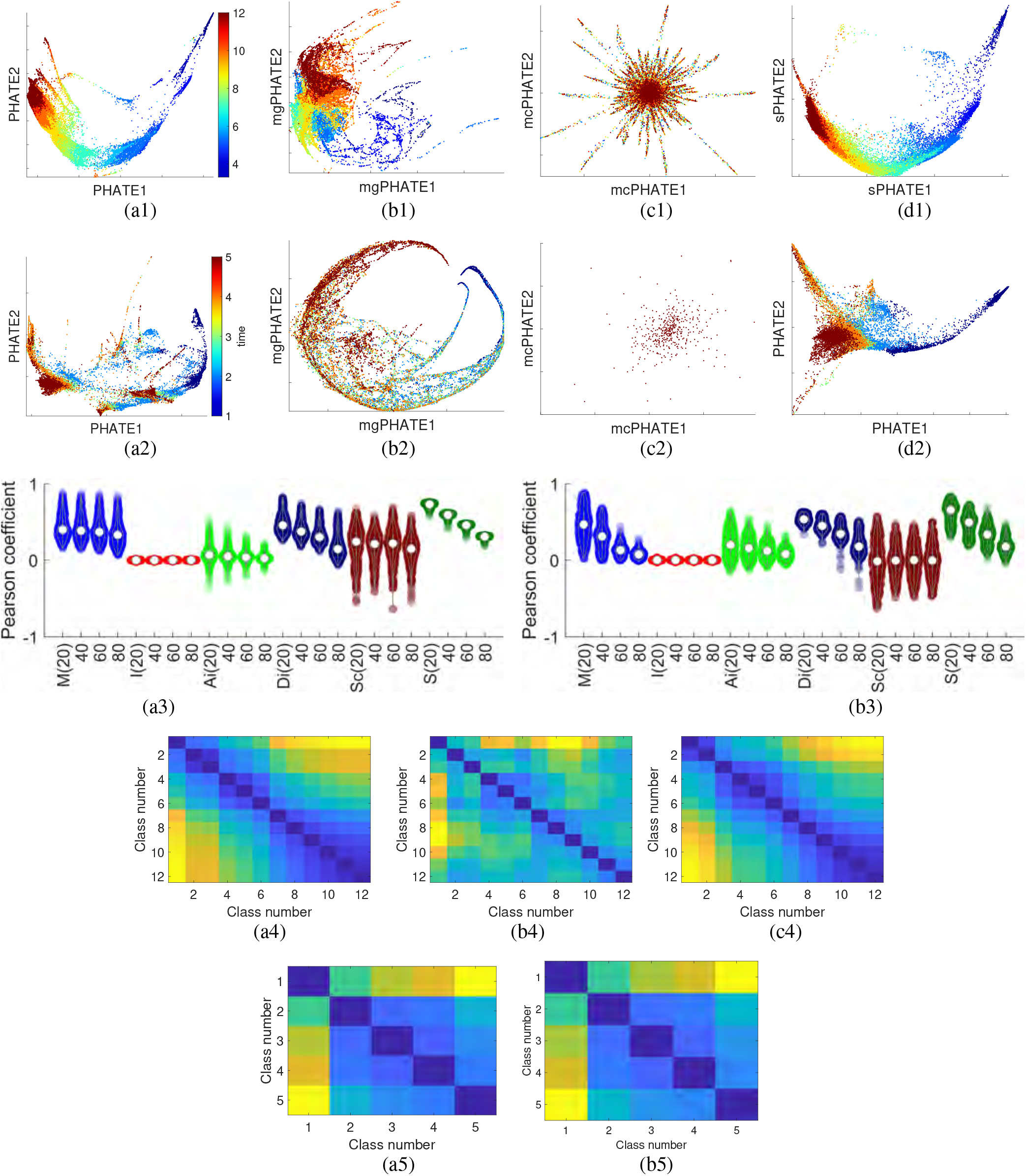
(preceding page). PHATE results from original data (a), imputed data from MAGIC (b), mcImpute (c) and SERM (d) for (1) zebrafish development data and (2) EB differentiation data. All the trajectories are better visualized in results from SERM imputation. MAGIC and mcImpute distort the data in both cases, whereas SERM keeps the consistency of the data intact in both cases. The colorbar for (a1-d1) denotes the hpf (hours post fertilization). The colorbar of (a2-d2) represents 1-(0-3 days), 2-(6-9 days), 3-(12-15 days), 4-(18-21 days) and 5-(24-27 days)). Pearson coefficient between the imputed data and ground truth data for all the methods are shown for zebrafish development data (a3), and EB differentiation data (b3), respectively. In (a3-b3), ‘M’ denotes MAGIC, ‘I’ denotes mcImpute, ‘AI’ denotes AutoImpute, ‘DI’ denotes DeepImpute, ‘Sc’ denotes SAUCIE and ‘S’ denotes SERM methods, respectively. The percent of randomly-zeroed expression values (20, 40, 60 and 80) in the ground truth data to create the noisy data are shown in braces. The spread of the violin denotes the variance of the coefficient values across different cells. The class to class geodesic distance across trajectory for zebrafish dataset are shown in (a4-c4) for original data, MAGIC-imputed data, and SERM-imputed data, respectively. The class to class geodesic distance across trajectory for EB differentiation dataset are shown in (a5-b5) for original data and SERM-imputed data, respectively.

The second dataset we use is scRNA-seq data from human Embryonic Stem (ES) cells differentiating as embryoid bodies (EBs)^35,37^. EB differentiation recapitulates key aspects of early embryogenesis and has been successfully used as the first step in differentiation protocols for certain types of neurons, astrocytes and oligodendrocytes, hematopoietic, endothelial and muscle cells, hepatocytes and pancreatic cells, and germ cells. Approximately 31,000 cells were measured, equally distributed over a 27-d differentiation time course. Samples were collected at 3-d intervals and pooled for measurement on the 10x Chromium platform. The PHATE embedding of the EB data, original and imputed with six methods, are shown in Fig. 4 (a2-d2) and Fig. S9 (a2-c2). Like with the zebrafish development data, most methods (MAGIC, mcImpute, AutoImpute) produce highly distorted results. PHATE results from SERM and DeepImpute-imputed data shows less distortion and the trajectory can be seen clearly.

The Pearson coefficients between the original data and imputed data by different methods are shown in Fig. 4 (a4, b4) for the above two datasets. Similar to cluster analysis datasets, SERM achieves the highest coefficients in most cases. MAGIC and DeepImpute perform comparatively better than other three techniques.

To demonstrate the efficiencies of all imputation methods in finding the correct trajectories quantitatively, the geodesic distances across the inter-class (each class corresponds to one timepoint) trajectories for the zebrafish development and EB datasets are computed, and the distance matrices are presented in fourth and fifth rows of Fig. 4 and Fig. S10, respectively. The distance matrices show distance increments as we move from a smaller to a larger number of classes, which is evident in the heatmaps of AuotImpute, DeepImpute and SERM-imputed data. MAGIC could not preserve the geodesic distances across the trajectory for the first dataset (Fig. 4 (b4)). It was not possible to compute the geodesic distance among data classes from mcImpute and SAUCIE-imputed data for both datasets and MAGIC-imputed data for second dataset because of the non-connectivity of the neighbors.

### SERM imputes big human cell landscape and mouse cell atlas data

The goal of this section is to demonstrate that SERM can be used to impute datasets of any size and complexity. To this end, we analyze two large scale datasets: the human cell landscape (HCL) and the mouse cell atlas (MCA). These two datasets have about 16,403 million data elements (599,926 cells × 27,341 genes) and 11,665 million data elements (333,778 cells × 34,947 genes), respectively, and no traditional method is able to impute these datasets within a reasonable time (days).

The HCL is a basic landscape of major human cell types created based on samples from a Han Chinese population using Microwell-seq technology^38^. Donated tissues were perfused or washed and prepared as single-cell suspensions using specific standard protocols. The analyses included samples of both fetal and adult tissue and covered 60 human tissue types (two to four replicates per tissue type, in general). Seven types of cell culture, including induced pluripotent stem (iPS) cells, embryoid body cells, hematopoietic cells derived from co-cultures of human H9 and mouse OP9 cells^39^, and pancreatic beta cells derived from H9 cells using a seven-stage protocol were also analyzed^40^. Single cells were processed using Microwell-seq^41^ and sequenced at around 3,000 reads per cell; data were then processed using published pipelines^8^. In total, 702,968 single cells passed the quality control tests (please see^38^ for details of the quality control tests). Following the original authors, we use 599,926 cells with 63 unique cell types and 59 unique tissue types for our analysis.

For creating the MCA, the same technology, Microwell-seq, is used^41^. Mammary gland (virgin, pregnant, lactation and involution), uterus, bladder, ovary, intestine, kidney, lung, testis, pancreas, liver, spleen, muscle, stomach, bone marrow, thymus, prostate, cKit+ bone marrow, bone marrow mesenchymal cells and peripheral blood samples from 6- to 10-week-old C57BL/6 mice were collected. Moreover, fetal liver, fetal lung, fetal stomach, fetal gonad, fetal brain, fetal intestine, fetal placenta and mesenchymal tissues, in addition to neonatal brain, neonatal skin, neonatal calvaria, neonatal rib and neonatal muscle samples, were collected. Tissues were carefully washed and prepared into single-cell suspensions with optimized protocols. Several cultured cells derived from mouse tissues: 3T3 cells, embryonic stem (ES) cells, trophoblast stem (TS) cells, and mesenchymal stem cells (MSCs) were also added to the analysis. Single cells were then sequenced with Microwell-seq. The sequencing data were processed using published pipelines^8^. > 400, 000 single cells from > 50 mouse tissues and cultures were analyzed in total. Following the original authors, we analyze 333,778 cells with 52 unique cell types and 47 unique tissue types.

The t-SNE visualization of the HCL after SERM imputation is shown in Fig. 5 (a), whereas the t-SNE visualization of the non-imputed data is shown in Fig. S11. From these figures, it is seen that the data classes are better clustered with SERM-imputed data due to the presence of more differentiating features among data classes. 19 new clusters can be found from the imputed data in comparison to the unimputed data. The data cluster numbers are shown in Figs. 5(a) and S11 (a) for both the imputed and unimputed data. In the original data, the maximum number of data clusters that can be found is 43, whereas in the SERM-imputed data, there are 62 clusters (out of 63 total clusters). Similarly, in the case of MCA, the total number of clusters found in the imputed case is 49 (out of 52) (Fig. 5(b)), whereas in unimputed data, there are 42 data clusters (Fig. S11 (b)). The accuracy of clusters in SERM-imputed data can be seen in the quantitative results shown in Fig. 6. All the indices (clustering accuracy, AR, Rand, Hubert and NMI) are higher in case of SERM-imputed data than the unimputed data. It should be noted that deep learning based methods (SAUCIE and DeepImpute) are able to analyze HCL and MCA datasets because of their scalability. However, we did not show the SAUCIE results because they did not show any data clusters and are in general significantly worse that the unimputed data. The DeepImpute data processing did not finish in two weeks for both datasets and were abandoned.

**Figure 5.**
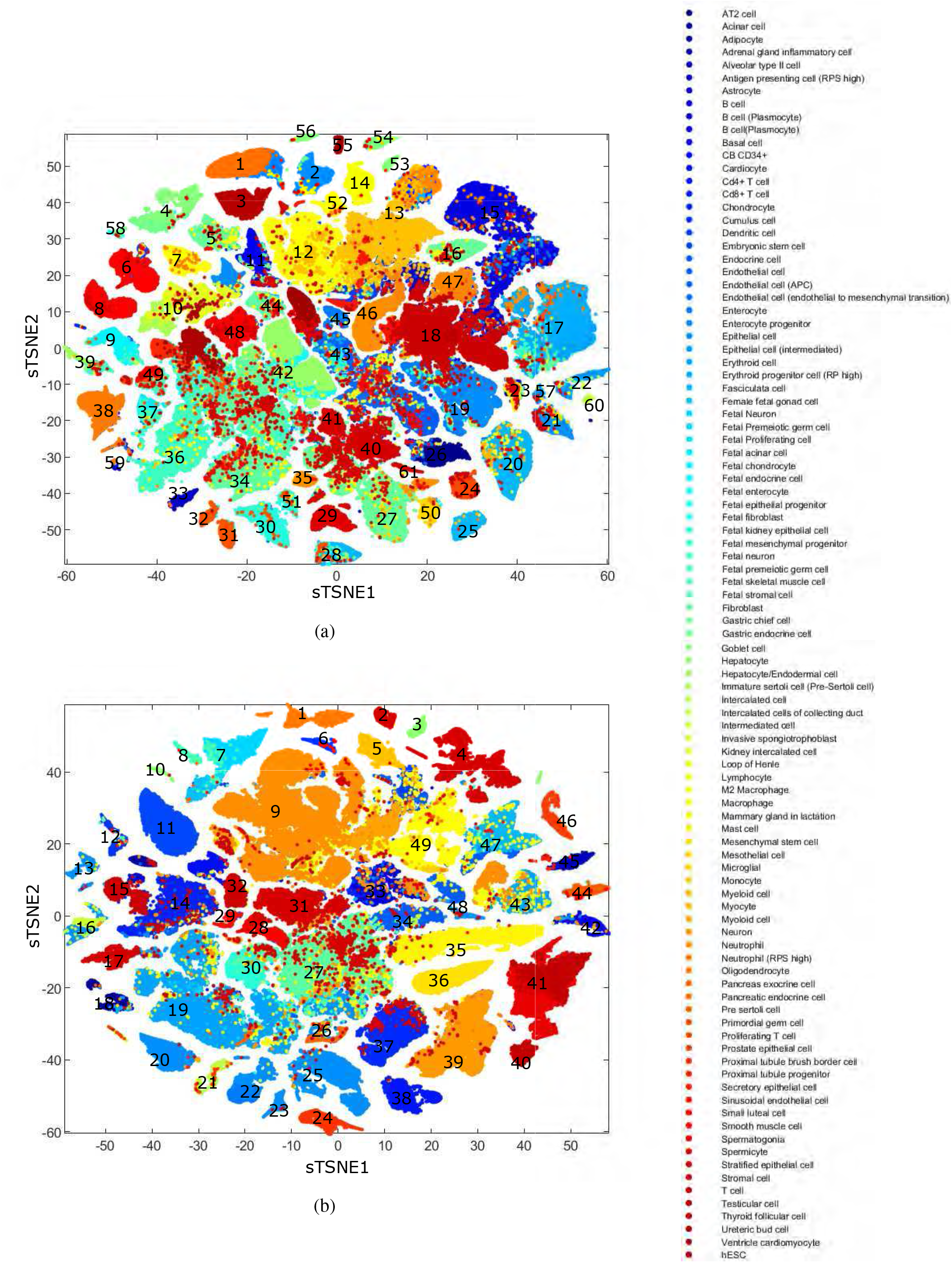
(preceding page). t-SNE visualization of human cell landscape (a) and mouse cell atlas (b) data imputed by SERM.

**Figure 6.**
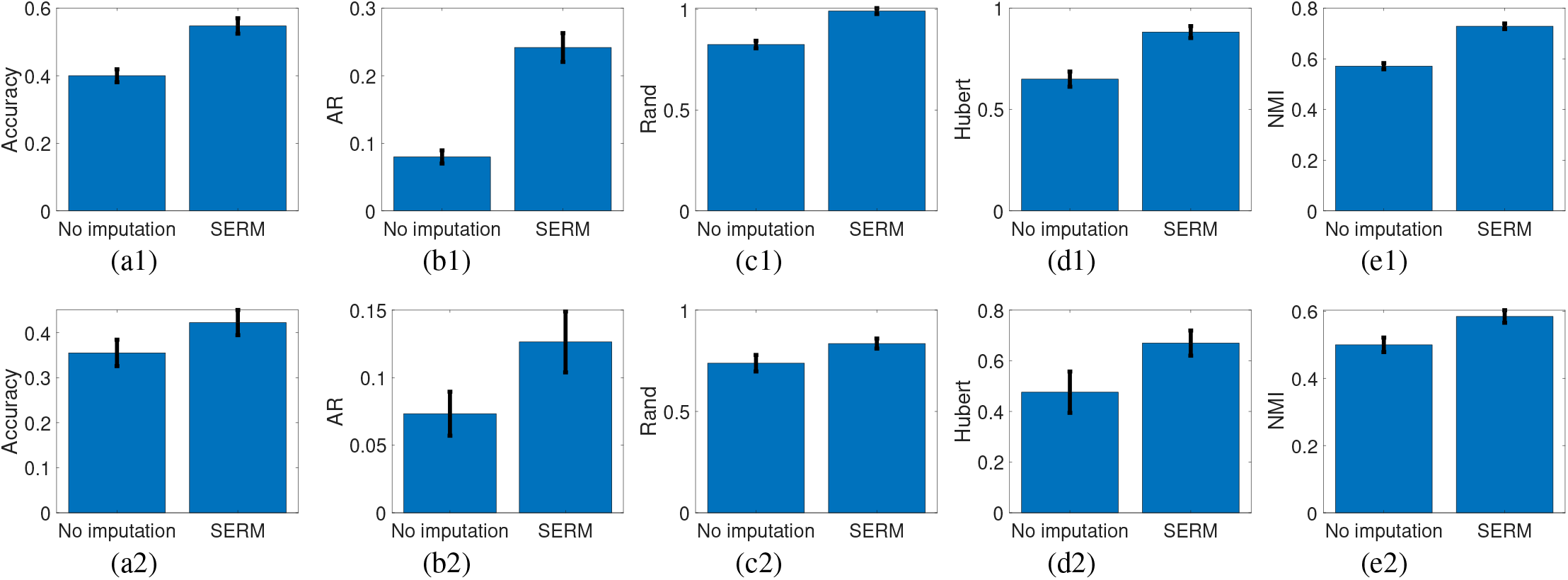
Accuracy (a), AR (b), Rand (c), Hubert (d) and NMI (e) indices of unimputed and SERM-imputed data for human cell landscape (1) and mouse cell atlas (2) datasets. Error bars represent the standard deviation of the indices for 1000 different initializations of k-means clustering.

More experiments are performed on single cell sequencing datasets from different established databases. PCA, t-SNE and UMAP visualizations of IDH-mutant gliomas data (section 4), single-cell data of pediatric midline gliomas (section 5), melanoma intra-tumor heterogeneity data (section 6), Div-Seq data (section 7), intestinal immune cell atlas (section 8), Drop-seq data (section 3), perturb-seq data (sec 10), mouse retina (sec 11), optimal transport analysis of iPSC reprogramming (sec 12) are included in Fig. 7 and the supplementary for SERM, mcImpute and MAGIC demonstrations. From these results, we observe that only in a few cases does performance of MAGIC and mcImpute compare to that of SERM. However, no other method can perform as well in all cases as consistently as SERM does.

**Figure 7.**
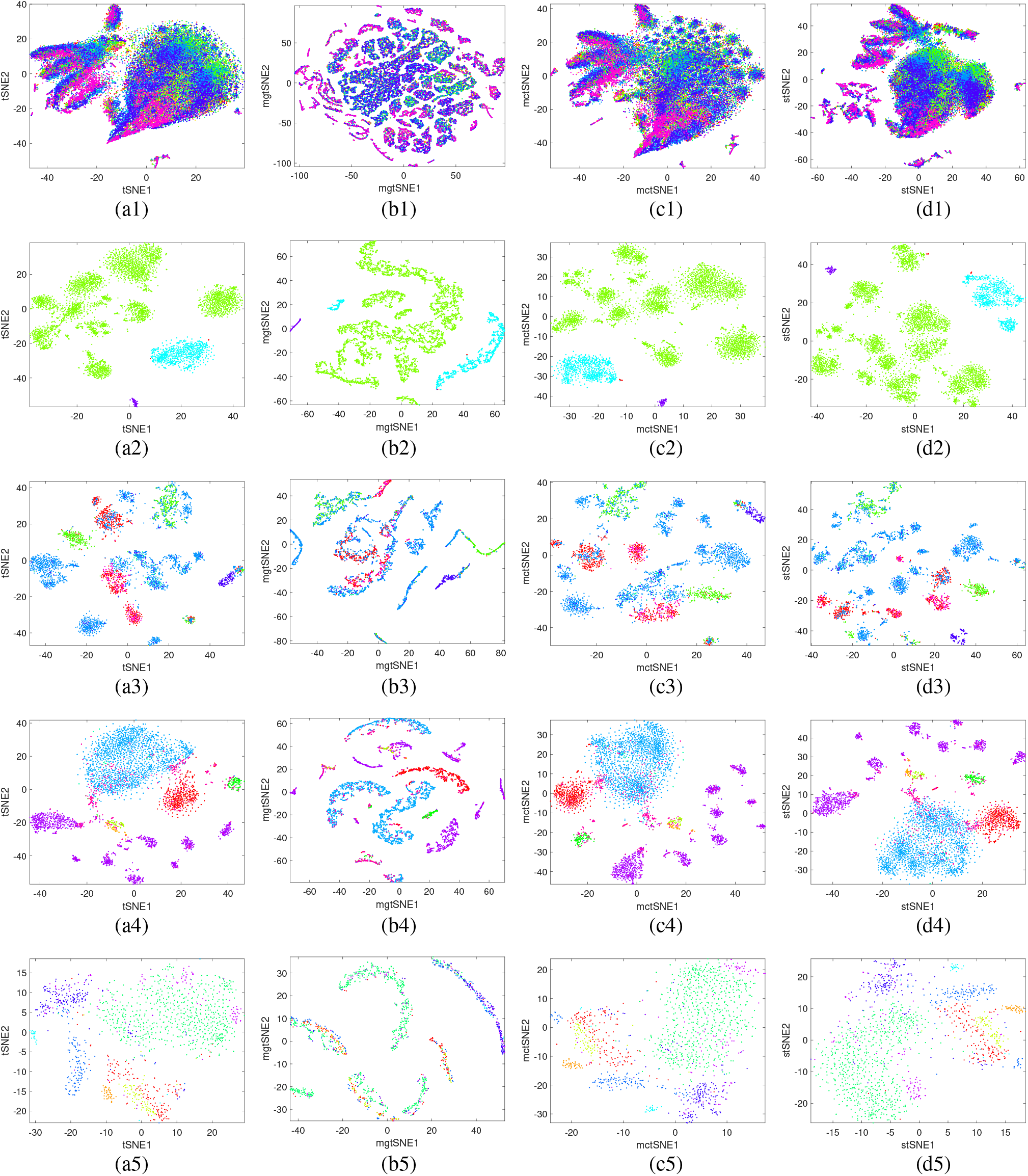
t-SNE results from original data (a), imputed data from MAGIC (b), mcImpute (c) and SERM (d) for (1) Drop-seq, (2) IDH-mutant gliomas, (3) pediatric midline gliomas, (4) melanoma intra-tumor heterogeneity, and (5) Div-Seq datasets. All the classes are better visualized in results from SERM imputation. MAGIC and mcImpute distort the data in many cases, whereas SERM keeps the consistency of the data intact in every cases. See methods for data description. Legends are added in the supplementary information.

It is also important to note that SERM is more computationally efficient than other state-of-the-art data imputation techniques. For example, SERM can analyze data with 0.6 million cells and 30,000 genes in less than 18 minutes on a personal computer with an Intel Core i9 processor and 64GB RAM, whereas MAGIC, mcImpute, AutoImpute and DeepImpute failed to analyze the same data within 3 days. The computational speeds of different techniques are reported for different numbers of data points in the supplementary (section 13). SERM is significantly much faster than all other methods. The underlying reason for this unprecedented enhancement in computational efficiency is that SERM uses a very fast histogram equalization operation to impute the data, so computationally intensive calculations such as determining the similarity between cells or genes are not required. SAUCIE is computationally more efficient than other methods except SERM.

## Discussion

Genomic sequencing techniques are now increasingly focused on the characterization of single cells. For many practical applications, such as data dimensionality reduction, visualization, and cellular spatial and temporal mapping from the single cell sequencing data, rectification of the expression values via reliable imputation is a prerequisite. In reality, however, the existing gene imputation methods suffer from a number of drawbacks: 1) a specific model for gene expression values is often assumed, which may not reflect the actual data distribution in many practical cases; 2) the data is imputed based on the similarity among a few cells or genes, which is susceptible to errors as the global relationship among cells and genes is not leveraged; 3) all zero expression values are often viewed as dropout events, which may not be true in practical settings; and 4) a priori information about the data is required to tune the hyper parameters of some methods. The proposed SERM mitigates these limitations through an effective learning strategy and significantly improves the imputation accuracy. It should be noted that SERM also assumes specific distributions for the expression data (such as exponential, Gaussian, Rayleigh). However, parameters of these distributions are learnt in SERM, which ensures the proper adaptation of the model to the data.

Data distortion, a condition when dropout positions in an expression matrix get filled with inaccurate values, is one of the main concerns in gene imputation. Preventing the distortion is critically needed in applications like trajectory mapping, where the accurate position of each cell is required to find the trajectory continuum. In our study, we found that most methods except SERM distorted the data in many cases and were unable to uncover the underlying biological information. With data-driven learning of the data distribution, SERM shows remarkable performance in imputing the gene expression values with high fidelity for trajectory inference, as revealed in our extensive experiments.

Reliability and consistency are two important features characterizing the performance of a gene imputation technique. When the dropout rate is high, all the methods except SERM do not perform well and their results hardly correlate with the actual expression values (Fig. 3). Thus in practical experiments, where the dropout rate is unknown, it may be risky to use these techniques. On the other hand, SERM-imputed data shows high correlation with the actual expression data even for situations with high dropout rates. Thus, SERM may be a reliable choice in practical gene expression analyses.

SERM is scalable to data size and dimensionality, making it possible to impute large data with high computational efficiency. This has been demonstrated by imputing two very large datasets (HCL and MCA). To the best of our knowledge, there are currently no methods other than SAUCIE, available to impute such large data within a reasonable computational time. In the visualizations of projected data in a low dimension, 19 new clusters were found in the case of HCL, and 7 new clusters in the case of MCA after SERM-imputation. Thus, using SERM, discovery of novel biological information such as cell or gene types becomes feasible from very large scRNA-seq data, which would otherwise be impossible.

We evaluated SERM on twenty different scRNA-seq datasets from different biological systems and measurement technologies. It is remarkable that, in all cases, SERM accurately recovers data clusters (Figs. 2, 3, 5), cell developmental trajectories, and state transitions (Fig. 4). The consistent superior performance of SERM across all these datasets demonstrate its versatility in applications across genetic engineering and computational biology. Generally speaking, the issue of gene expression imputation here is a special case of missing data or missing values problem, which can occur in almost all imaging and measurement modalities. In practice, the strategy of dealing with the problem can significantly affect the conclusions to be drawn from the data^42–44^. Since SERM is based on the learning from the data, it is not restricted to a specific type of data. The method can adapt to a new data easily and may provide a broadly applicable strategy for missing data recovery across different disciplines. In addition to the missing value imputation, we emphasize that the proposed SERM also provides an effective solution in denoising the data, as demonstrated in Figs. 2–5. To certain extent, the proposed SERM method in dealing with tabulated data here is analogous to the commonly used data-driven super-resolution imaging techniques for imaging data^45^.

In our analyses, we found out that MAGIC distorts the t-SNE/UMAP visualizations of the data in many cases. This may be because of the smoothing effect from MAGIC algorithm and may not always imply the bad quality of the imputation. Rather, we found that MAGIC performs better than most techniques in terms of quantitative indices such as clustering accuracy, NMI and cluster quality indices. This finding is corroborated by an earlier study published by Hou et al^15^. Interestingly, SERM also has an smoothing effect on the data because of the histogram equalization. However, we did not find any distortion in visualization from SERM-imputed data.

Despite its success, SERM is not without limitations. In particular, SERM learns the gene expression model from a small fraction of the data. This poses a challenge in dealing with small genomic data-sets (say a few single cells of *n* < 20), as the estimated distribution in SERM may diverge from the actual data distribution, leading to inaccurate imputation values.

In summary, we have proposed a novel computational strategy for gene expression recovery. The technique can impute gene expression data of different levels of dropout rate with unprecedented accuracy, reliability, and consistency. Going beyond traditional imputation techniques, SERM learns the data distribution and applies the knowledge to the subsequent calculation under the condition of self-consistency. SERM is computationally efficient and scalable to large data sizes. As SERM is data-driven and can adjust to any data type, we envision that SERM will play an important role in various biomedical and biocomputational applications.

## Methods

The success of SERM is attributed to its adaptability and scalability. The SERM strategy involves the following major steps:

1. (Modeling-Fig. 1 steps 1-3) The goal of this step is to determine the distribution model that best represents the gene expression data. First, a subset of the full original dataset must be defined. This data subset is transformed into an ideal version of itself via an unsupervised deep learning technique that performs compression and decompression of the data. This process removes noise and retains the essential latent features of the data, thus outputting an ideal and denoised representation of the original data. A histogram of the ideal expression values is computed, and distribution fitting is performed on the histogram to output what will be referred to as the learned pdf in the following steps.
2. (Imputation-Fig. 1 steps 4-6) In this step, the values in the gene expression matrix are systematically imputed based on a sliding window. A rectangular ROI is defined with width and height smaller than the width and height of the full expression matrix. Depending on its position, the ROI serves as a window into the matrix where only the values within the window are manipulated in a single instance. The sliding parameters must then be defined to determine the distance that the window shifts for each iteration. The window shifts in both the x- and y-directions, but the amount of x-direction sliding must be less than or equal to the window width, and the amount of y-direction sliding must be less than or equal to the window height. The sliding distance should be divisible by the size of the matrix. The number of iterations is determined by the number of times the window must slide in order to incorporate the full matrix. In each iteration, a histogram of the values in the window is computed. This histogram is equalized to the histogram of the ideal values found in the previous step using the learned pdf. The new window values found through histogram equalization substitute for the original values. The window continues sliding until the entire matrix is imputed.
3. (Consistency-Fig. 1 step 7) As previously stated, the distance that the window slides must be smaller than or equal to the size of the window. This ensures global coverage of the matrix – no values are skipped. If the sliding distance is strictly smaller than the window size, then the values in each window overlap with those in the adjacent windows. Cross-window consistency is built-in for this choice of parameters. In the case that the sliding distance is exactly equal to the window size, adjacent windows do not overlap, and the values in each window are imputed independently from the rest of the matrix. Bilinear resampling is performed to impose global consistency across all the independent windows.

The detailed theory behind each step of SERM is discussed below.

### Learning data distribution

Deep neural networks (DNNs) have shown promising results in the classification of biomedical data through learning complex structures in the data across many domains through training with example datasets^46^. Autoencoders^47^ (consists of an encoder and a decoder) are a class of DNNs where no labeled data are necessary. This network learns to compress the high-dimensional data by adjusting the weights of its neurons in an unsupervised manner. It learns only the essential latent features ignoring non-essential sources of variation such as random noise^23^. Therefore, the compressed latent data of network reflects the high dimensional ambient data space in lower dimensionality and captures the underlying true data manifold. Thus, when the autoencoder reconstructs the high-dimensional data, it represents the ideal version of the high-dimensional denoised data^23^. We use a subset (*S_o_* of size *q_s_ × m_s_*) of the expression data (*S* of size *q × m*) as input to the network and learns the distribution of the ideal expression data.

#### Deep learning optimization

The deep learning architecture used in SERM consists of two parts, an encoder and a decoder. The encoder stage the network takes the input *x_d_* ∈ ℝ^*Q*^ (*x_d_* is a row of *S_s_*) and maps it to a latent space **h** ∈ ℝ^*P*^, where *Q* > *P* and **h** is defined as

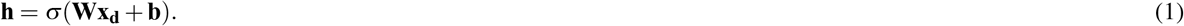

Here, σ is an element-wise activation function which can be a sigmoid or linear function, **W** is a weight matrix and **b** is a bias vector. Weights and biases are usually initialized randomly, and then updated iteratively during training using backpropagation technique. Subsequently, the decoder stage of the network maps the latent variable **h** to the reconstruction data 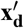. This process can be written as

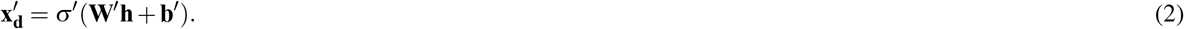

While training the network in SERM, reconstruction error (mean squared error) is minimized. The mean squared error (also called loss) can be expressed as

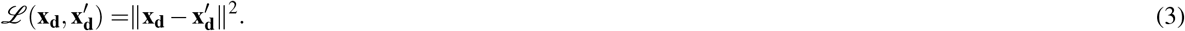

After we replace 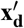 in equation 3 using equation 2, we obtain

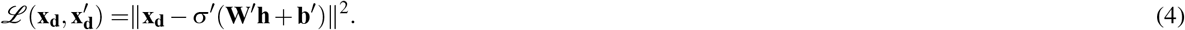

If we replace **h** in equation 4 using equation 1, we obtain the following expression for the loss function:

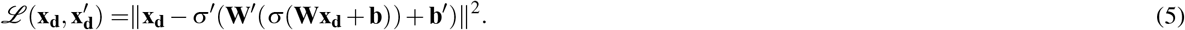

The resulting latent space **h** is the desired dimensionality-reduced data. To improve the performance of the network, two additional terms (*L*_2_ regularization and sparsity regularization terms) are generally added to the loss function as follows:

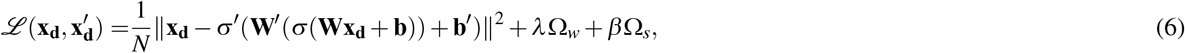

where *N* is the total number of training examples and λ and β are coefficients of *L*_2_ regularization and sparsity regularization terms. The sparsity regularization term Ω_*s*_ is defined as

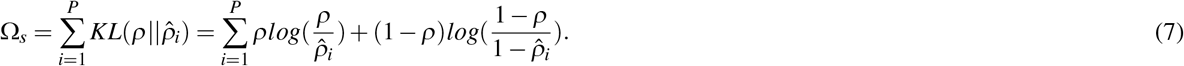

Here *ρ* is the sparsity proportion parameter denoting the desired value of the average activation value of the neurons in the network. Average activation value of the *i*-th neuron is computed as

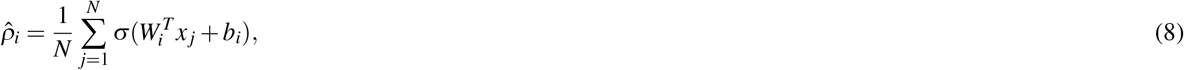

where *x _j_* is the *j*-th training example. In equation 6, Ω_*w*_ denotes the *L*_2_ regularization term and is defined as

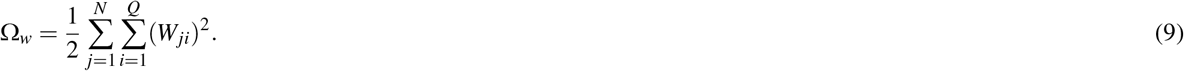

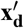 are the rows of reconstructed output matrix *S_o_*, which is of the same size as *S_s_*.

#### Histogram computation

Consider the gene expression matrix 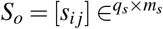, where *q_s_* is the number of cells and *m_s_* is the number of genes. For each expression value, the number of elements with that value in *S_o_* is counted. The collection of these counts for all the expression values is referred to as the histogram of that region. This function is in general an estimate of the expression density function. Let 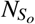 and 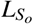 denotes the number of matrix elements and possible expression values in *S_o_*. Furthermore, we define the normalized histogram associated with the possible gene expression values in *S_o_* as follows

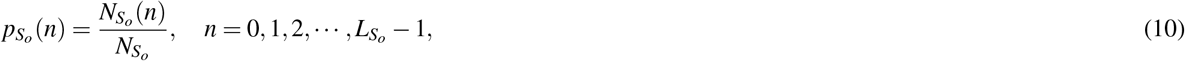

where 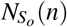 is the number of elements in the tile with the gene count *n* such that 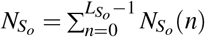.

#### Curve fitting

We fit curves with the following equations to 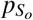.

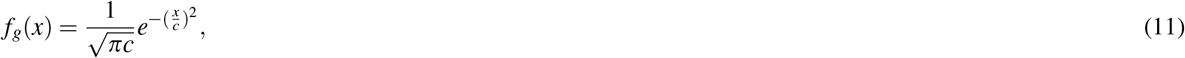

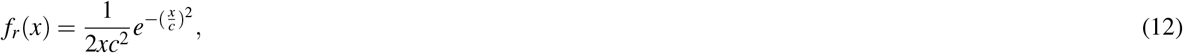

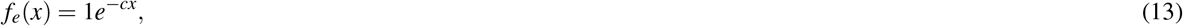

where *f_g_*, *f_r_* and *f_e_* denote Gaussian, Rayleigh and exponential probability distribution function (pdf). The pdf with parameter *c* that results in minimum root mean square (RMSE) is chosen for next steps of SERM.

### ROI selection and histogram equalization

Let us denote an ROI of the expression matrix *S* as *U* (*i*, *j*), where *i* = 1,…, *R_y_* and *j* = 1,…, *R_x_*. If the ROI is shifted by *R_l_* row and *R_m_* column in each change of *i* and *j*, the (*c*, *d*)th ROI is defined as *U_c_*_,*d*_ = *U* (*i* + *c* * *R_l_*, *j* + *d* * *R_m_*). The expression matrix is zero-padded symmetrically by adding 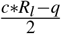 and 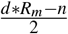 at starting and ending of the expression matrix. Here *c* and *d* are the nearest greater integer resulting from 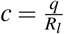 and 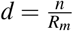.

Consider the gene expression matrix *S* = [*s_i j_*] ∈^*q*×*m*^, where *n* is the number of cells and *m* is the number of genes. We divide the expression matrix *S* is divided into *N* ≥ 4 submatrices (tiles), which we denote by *S_i_*, *i* = 1, 2, · · ·, *N*. Let 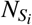 and 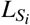 denotes the number of matrix elements and possible expression values, in each tile *S_i_*, respectively. Furthermore, we define the normalized histogram associated with the possible gene expression values in the tile *S_i_* as follows

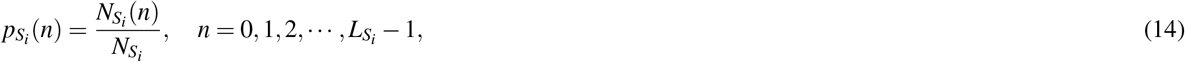

where 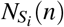 is the number of elements in the tile with the gene count *n* such that 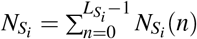. The corresponding cumulative distribution function (CDF), is given by

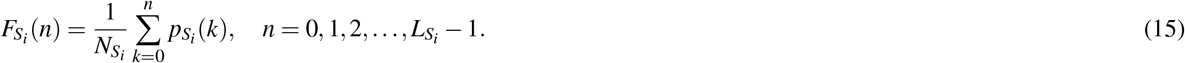

#### Adaptive equalization of histogram

The goal of the histogram adaptive equalization is to find a transformation for the expression values such that the transformed CDF is approximated by the CDF of *f* (*x*), where *f* (*x*) can be *f_g_*(*x*), *f_r_*(*x*) or *f_e_*(*x*). As example, the CDF of an exponential distribution can be written as

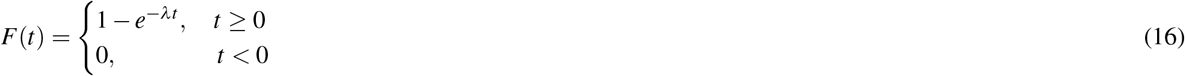

where λ denotes the decay parameter. SERM uses the transformation 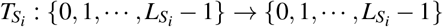 for each tile to map the expression values to their new values. In particular, consider transforming the expression counts by

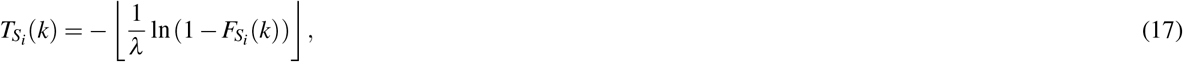

where the floor function ⌊*x*⌋ is the biggest integer not exceeding *x*. The map 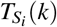 is then applied to each expression value in the tile *S_i_*.

The main intuition behind this transformation comes from the inverse sampling method. In particular, let *X* denotes continuous random variables whose CDF is strictly increasing on the possible value on 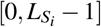 with densities *p_X_* and *p_Y_*, respectively. Consider the transformation

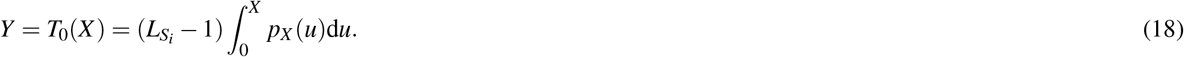

Then, *Y* has the uniform distribution, *i.e*., 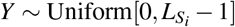. To see this note that

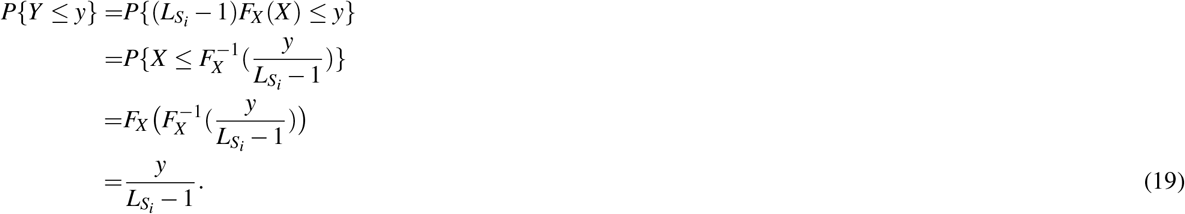

Furthermore, using the inverse transform sampling technique, the random variable

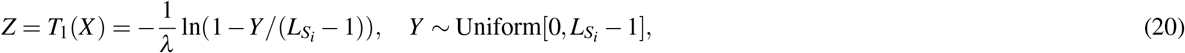

has the exponential distribution, *i.e*., *Z* ~ *F*(*t*). Combining the transformations *T*_0_ and *T*_1_, we conclude that

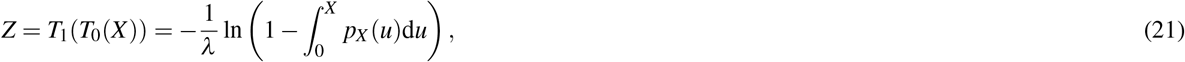

has the desired exponential distribution. Now, the transformation 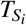 defined in Eq. (17) is the discretized version of the continous transformation *T*_1_ ° *T*_0_.

### Sliding ROI

The ROI is slided throughout the whole expression matrix and gene imputation is performed at each ROI to obtain the resulting imputed gene expression matrix from SERM.

### Resampling for patch consistency

In the previous step, the imputation is performed on each tile by matching the transformed clipped histogram of each time with an exponential distribution. We now incorporate the histograms of different tiles into a consistent framework by applying a bilinear resampling. Let 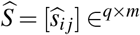 denotes the gene expression matrix after the imputation of each tile via the histogram clipping and equalization step. We subsequently resample each expression value 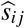 using the values of lateral elements 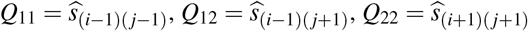, and 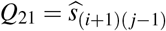. In particular, let 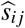 denotes the resampled value which is computed by the following sample average

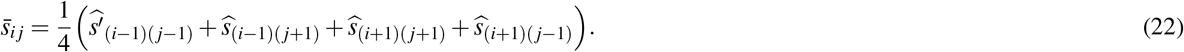

For the boundary elements of the matrix 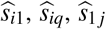, and 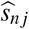 the above formula is used in conjunction with the zero padding of the matrix 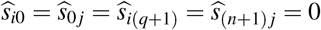 for all *i* ∈ {1,2, …, *q*} and *j* ∈ {1,2, …, *m*}. The interpolation formula in Eq. (22) can be de rvied via th e discretiz ation of the bilinear interpolation of continuous functions of two variables. The resulting matrix 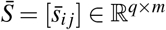 from resampling in Eq. (22) yields the desired imputed matrix.

### Simulation procedures

All the simulated data were created in R with Splatter simulator.

### Implementation and parameter settings

MATLAB (MathWorks Inc., Natick, MA, USA) is used to implement the SERM technique. In the case of the deep learning optimization of SERM, a linear function has been used as the decoder transfer function and a logistic sigmoid function as the encoder transfer function. Mean squared error between the input data and reconstructed data from the decoder is used as the loss function of the network. The maximum number of epochs is set to be 20. The *L*_2_ regularization parameter (λ) is chosen as as 0.001 and the sparsity proportion parameter (*ρ*) as 0.05. The sparsity regularization parameter (*β*) is chosen as 1.6. Among these parameters, loss function, encoder transfer function, *L*_2_ regularization parameter, sparsity proportion parameter and sparsity regularization parameter are set to the default MATLAB parameters set by Matlab. Setting the encoder transfer function to a sigmoid function and the decoder transfer function to a purely linear one is an ideal setting for better performance as described by Vincent et al.^47^. 20 epochs is enough for reaching the global minimum in all our analyses. However, these parameters may be tuned for better performance of SERM in specific applications.

For curve fitting, the ‘Trust-Region’ algorithm is used with mean square error as the cost function. The starting point, lower limit and upper limits of *P* for curve fitting to all equations (*f_g_*, *f_r_*, *f_e_*) were 1, 0 and infinity. For Gaussian and Rayleigh equation fitting, the starting point, lower limit and upper limits of *c* were 0.15, 0.1 and 0.3, respectively. For exponential equation fitting, the starting point, lower limit and upper limits of *c* were 5, 5 and 20, respectively. These values were chosen based on fitting the equations to ideal data generated using the Splatter simulator. In SERM, the window length is set to one-fourth of the matrix length, and the window height is set to one-fourth of the matrix height (*N* = 4(2×2)). The x- and y-sliding parameters are set to the length and height of the window. t-SNE implemented by MATLAB is have been used to produce the t-SNE visualizations of this method. UMAP is implementation from by the authors and used has been used with default parameters^48^.

### Datasets

#### Cellular taxonomy of the mouse bone marrow stroma

In this dataset, single-cell RNA sequencing (RNA-seq) was used to define a cellular taxonomy of the mouse bone marrow stroma and its perturbation by malignancy^31^. Seventeen stromal subsets were identified expressing distinct hematopoietic regulatory genes spanning new fibroblastic and osteoblastic subpopulations including distinct osteoblast differentiation trajectories. Emerging acute myeloid leukemia impaired mesenchymal osteogenic differentiation and reduced regulatory molecules necessary for normal hematopoiesis. These data suggest that tissue stroma responds to malignant cells by disadvantaging normal parenchymal cells. This taxonomy of the stromal compartment provides a comprehensive bone marrow cell census and experimental support for cancer cell crosstalk with specific stromal elements to impair normal tissue function and thereby enable emergent cancer.

#### sNucDrop-seq data

The next dataset we analyze is from a massively parallel scRNA-seq technology namely sNucDrop-seq (single-nucleus RNA-seq approach), which is free of enzymatic dissociation and nucleus sorting^32^. This technology allows unbiased isolation of intact single cells from complex tissues such as adult mammalian brains, which is a daunting task. The authors profiled 18,194 nuclei isolated from cortical tissues of adult mice. The authors demonstrated through extensive validation that sNucDrop-seq not only accurately reveals neuronal and non-neuronal subtype composition with high accuracy but also allows in-depth analysis of transient transcriptional states driven by neuronal activity.

#### Mouse intestinal epithelium

Intestinal epithelial cells absorb nutrients, respond to microbes, function as a barrier and help to coordinate immune responses. 53,193 individual epithelial cells from the small intestine and organoids of mice were profiled, which enabled the identification and characterization of previously unknown subtypes of intestinal epithelial cell and their gene signatures. We analyzed 7,216 cell data using SERM and other imputation methods and reported the results.

#### Human engineered neural cells

The dataset analyzed in this section is from a study on human engineered neural tissues^34^. Single cell RNA-seq data of human induced neuronal cells cultured in two different conditions either with mouse astrocytes or with differentiated human astrocytic cells are. Here we present only data from human cells. Two protocols were followed for data acquisition:

Lane-1: Human embryonic stem cell (hESC) derived induced neuronal cells and mouse astrocytes were co-cultured at 1:1 ratio in a 3D Composite Hydrogel with 4X CRS (12.5 mM CaCl2) at 30 million cells per ml.
Lane-2: hESC derived induced neuronal cells and human astrocytic cells differentiated from hESCs were co-cultured at 1:1 ratio in a 3D Composite Hydrogel 4X CRS at 30 million cells per ml.

#### Zebrafish embryogenesis

We use here a large dataset obtained through profiling 38,731 cells from 694 embryos across 12 closely separated stages of early zebrafish development^36^ using a massively parallel scRNA-seq technology named Drop-seq^2^. Data were acquired from the high blastula stage (3.3 hours postfertilization (hpf), moment after transcription starts from the zygotic genome) to six-somite stage (12 hours after postfertilization, just after gastrulation). Most cells are pluripotent at high blastula stage, where as many cells have differentiated into specific cell types at the six-somite stage.

#### Analysis of scRNA-seq data from Drop-seq technology

Cells, the basic units of biological structure and function, vary broadly in type and state. Single cell genomics can characterize cell identity and function, but limitations of ease and scale have prevented its broad application^2^. Drop-seq is a strategy for quickly profiling thousands of individual cells by separating them into nanoliter-sized aqueous droplets, associating a different barcode with each cell’s RNAs, and sequencing them all together. Drop-seq analyzes mRNA transcripts from thousands of individual cells simultaneously while remembering transcripts’ cell of origin. In Ref.^2^, the authors analyzed transcriptomes from 44,808 mouse retinal cells and identified 39 transcriptionally distinct cell populations, creating a molecular atlas of gene expression for known retinal cell classes and novel candidate cell subtypes. Drop-seq will accelerate biological discovery by enabling routine transcriptional profiling at single cell resolution.

#### IDH-mutant gliomas

Tumor subclasses differ according to the genotypes and phenotypes of malignant cells as well as the composition of the tumor microenvironment (TME)^49^. The authors dissected these influences in isocitrate dehydrogenase (IDH)–mutant gliomas by combining 14,226 single-cell RNA sequencing (RNA-seq) profiles from 16 patient samples with bulk RNA-seq profiles from 165 patient samples. Differences in bulk profiles between IDH-mutant astrocytoma and oligodendroglioma can be primarily explained by distinct TME and signature genetic events, whereas both tumor types share similar developmental hierarchies and lineages of glial differentiation.

#### Single-cell analysis in pediatric midline gliomas

Gliomas with histone H3 lysine27-to-methionine mutations (H3K27M-glioma) arise primarily in the midline of the central nervous system of young children, suggesting a cooperation between genetics and cellular context in tumorigenesis^50^. While the genetics of H3K27M-glioma are well-characterized, their cellular architecture remains uncharted. In this dataset, the authors performed single-cell RNA-seq in 3,321 cells from six primary H3K27M-glioma and matched models. This study characterizes oncogenic and developmental programs in H3K27M-glioma at single-cell resolution and across genetic subclones, suggesting potential therapeutic targets in this disease.

#### Melanoma intra-tumor heterogeneity

To explore the distinct genotypic and phenotypic states of melanoma tumors, the authors applied single-cell RNA sequencing (RNA-seq) to 4645 single cells isolated from 19 patients, profiling malignant, immune, stromal, and endothelial cells^51^. Malignant cells within the same tumor displayed transcriptional heterogeneity associated with the cell cycle, spatial context, and a drug-resistance program. In particular, all tumors harbored malignant cells from two distinct transcriptional cell states, such that tumors characterized by high levels of the MITF transcription factor also contained cells with low MITF and elevated levels of the AXL kinase. Single-cell analyses suggested distinct tumor microenvironmental patterns, including cell-to-cell interactions. Analysis of tumor-infiltrating T cells revealed exhaustion programs, their connection to T cell activation and clonal expansion, and their variability across patients. Overall, in this dataset, the cellular ecosystem of tumors began to unravel and how single-cell genomics offers insights with implications for both targeted and immune therapies.

#### Div-Seq data analysis

Single cell RNA-seq provides rich information about cell types and states^52^. However, it is difficult to capture rare dynamic processes, such as adult neurogenesis, because isolation of rare neurons from adult tissue is challenging and markers for each phase are limited. Div-Seq technique is developed to solve this problem, which combines scalable single-nucleus RNA-Seq (sNuc-Seq) with pulse labeling of proliferating cells by EdU to profile individual dividing cells. sNuc-Seq and Div-Seq can sensitively identify closely related hippocampal cell types and track transcriptional dynamics of newborn neurons within the adult hippocampal neurogenic niche, respectively. This study contains the sNuc-Seq analysis performed as a part of the Div-Seq method development.

Using sNuc-Seq, the authors analyzed 1,367 single nuclei from hippocampal anatomical sub-regions from adult mice, including enrichment of genetically-tagged lowly abundant GABAergic neurons. sNuc-Seq robustly generated high quality data across animal age groups (including 2 years old mice), detecting 5,100 expressed genes per nucleus on average, with comparable complexity to single neuron RNA-Seq from young mice. Analysis of sNuc-Seq data revealed distinct nuclei clusters corresponding to known cell types and anatomical distinctions in the hippocampus.

### Competing methods

#### MAGIC

MAGIC was downloaded from https://github.com/pkathail/magic. MAGIC was run using default parameters specified as 20 for the numbers of principal components, 6 for the parameter t for the power of the Markov affinity matrix, 30 for the number of nearest neighbors, 10 for the autotune parameter and 99th percentile to use for scaling.

#### mcImpute, DeepImpute, SAUCIE and AutoImpute

mcImpute, DeepImputeS, AUCIE and AutoImpute were downloaded from https://github.com/aanchalMongia/McImpute_scRNAseq, https://github.com/lanagarmire/deepimpute, https://github.com/KrishnaswamyLa and https://github.com/kearnz/autoimpute. All these methods were run with default configurations.

### Computation of Pearson coefficient

Let us assume that *R_f_* and *R_e_* are the gene expression from dropout-free data and imputed data using different techniques. Let us assume, A is the dropout-free gene expression data for a single cell from *R_f_*, whereas B is the gene expression vector for the same cell from *R_e_*. Then the Pearson correlation coefficient between A and B is defined as

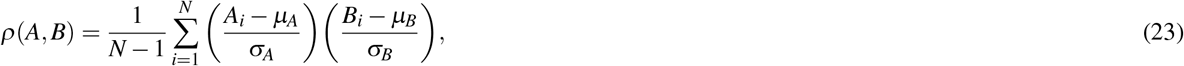

where μ_*A*_ and σ_*A*_ are the mean and standard deviation of A, respectively, and μ_*B*_ and σ_*B*_ are the mean and standard deviation of B.

### Computation of NMI, accuracy and cluster quality indices

We at first cluster the data into *N_g_* classes (*N_g_* is number of classes in ground truth label) by k-means clustering technique with Euclidean distance. We then find the best map of cluster labels in comparison to the ground truth labels. These cluster labels are then used to compute the NMI, accuracy and cluster quality indices: adjusted Rand (AR), Rand and Hubert. NMI is the normalized mutual information^53^ between the estimated labels and true labels computed following the work of Becht et al.^54^. Accuracy is the number of correctly found class labels divided by total number of class labels. Rand and AR index are computed using the formula reported in the works of Rand^55^ and Hubert et al.^56^.

## Data and code availability

The datasets generated during and/or analyzed during the current study are available within the manuscript and supplementary. Implementation codes are available in https://github.com/xinglab-ai/self-consistent-expression-recovery-machine.

## Acknowledgments

This work was partially supported by NIH (1R01 CA223667 and R01CA227713) and a Faculty Research Award from Google Inc.

## Author contributions statement

L.X. conceived the experiment(s), M.T.I conducted the experiment(s), M.T.I., X.L., M.B.K., J.Y.W., L.W., L.S., H.R. and W.Z. analyzed the results. All authors reviewed the manuscript.

